# Does co-expression of *Yarrowia lipolytica* genes encoding Yas1p, Yas2p and Yas3p make a potential alkane-responsive biosensor in *Saccharomyces cerevisiae*?

**DOI:** 10.1101/2020.09.16.299479

**Authors:** Yasaman Dabirian, Christos Skrekas, Florian David, Verena Siewers

## Abstract

Alkane-based biofuels are desirable to produce at a commercial scale as these have properties similar to our current petroleum-derived transportation fuels. Rationally engineering microorganisms to produce a desirable compound, such as alkanes, is, however, challenging. Metabolic engineers are therefore increasingly implementing evolutionary engineering approaches combined with high-throughput screening tools, including metabolite biosensors, to identify productive targets. Engineering *Saccharomyces cerevisiae* to produce alkanes can be facilitated by using an alkane-responsive biosensor, which can potentially be developed from the native alkane-sensing system in *Yarrowia lipolytica*, a well-known alkane-assimilating yeast. This putative alkane-sensing system is, at least, based on three different transcription factors (TFs) named Yas1p, Yas2p and Yas3p. Although this system is not fully elucidated in *Y. lipolytica*, we were interested in evaluating the possibility of translating this system into an alkane-responsive biosensor in *S. cerevisiae*. We evaluated the alkane-sensing system in *S. cerevisiae* by developing one sensor based on the native *Y. lipolytica ALK1* promoter and one sensor based on the native *S. cerevisiae CYC1* promoter. In both systems, we found that the TFs Yas1p, Yas2p and Yas3p do not seem to act in the same way as these have been reported to do in their native host. Additional analysis of the TFs suggests that more knowledge regarding their mechanism is needed before a potential alkane-responsive sensor based on the *Y. lipolytica* system can be established in *S. cerevisiae*.

## INTRODUCTION

Replacing petroleum-derived compounds with more environmentally friendly alternatives is needed due to the increasing global climate changes^*1*^. One approach to meet this need is the use of engineered microorganisms to produce industrially relevant products from bio-based renewable substrates^*2*^. However, engineering microorganisms to fit the industrial needs in terms of product spectrum and process economy is challenging and laborious due to our limited understanding of metabolism^*3*^. This challenge is exemplified by the long-standing desire to commercially produce 3^rd^ generation biofuels, also known as advanced biofuels, which are produced from non-food lignocellulosic biomass and can replace fuels used for heavy duty vehicles and ships as well as jet fuels^*4*^. Despite the extensive research in this area, production of advanced biofuels, such as alkanes, using engineered microorganisms has not yet reached commercial scale.

Rational engineering strategies, such as various DNA assembly methods^*5, 6*^ and efficient integration of complex pathways using, for example, CRISPR (clustered regularly interspaced short palindromic repeats) editing^*7*^, have advanced and facilitated cell factory development^*2*^. However, such approaches are still often trial-and-error processes due to the inherently complex and dynamic host metabolism. To overcome some of the challenges encountered in rational engineering, researchers are increasingly implementing approaches inspired from natural evolution processes^*8*^, such as evolutionary engineering and directed evolution. The use of these methods can help identify desirable engineering targets as well as overcoming the often-encountered inefficiency of naturally occurring enzymes.

In evolutionary engineering, strains with beneficial mutations are selected through survival under selective pressure, and such mutations can either arise naturally or be artificially imposed by the researcher. Directed evolution is commonly used for creating a large number of different variants of a gene encoding, for example, a specific enzyme to find variants with improved properties^*9*^. To identify promising strains or enzyme variants from either evolutionary engineering or directed evolution approaches, high-throughput screening methods are necessary^*10*^. Metabolite biosensors, for example transcription factor (TF)-based biosensors, are promising tools for high-throughput screening as these can provide fast and semi-quantitative measurements of a compound of interest^*11, 12*^.

The aim of this study was to develop an alkane-responsive, TF-based, biosensor to facilitate the development of an alkane overproducing yeast strain. Previous studies successfully produced alkanes through rational engineering of *Saccharomyces cerevisiae* using different biochemical pathways^*13, 14*^. Despite these achievements, the level of alkanes produced is so far low and far from being industrially relevant. One challenge with regard to alkane production concerns the engineering of the low-activity enzyme aldehyde-deformylating oxygenase (ADO)^*15*^, which converts fatty aldehydes to alkanes/alkenes, to obtain improved kinetic properties. There is therefore an interest in implementing directed evolution approaches to create variants of the ADO enzyme with improved properties such that improved alkane titers can be achieved. In order to identify appropriate variants among a large library of enzymes, an alkane-responsive biosensor is necessary^*16*^.

An alkane-responsive sensor based on a bacterial transcriptional activator system has previously been developed in *Escherichia coli* for the purpose of identifying promising microbial cell factories producing alkanes^*17*^. As it is commonly challenging to implement prokaryotic activators in a eukaryotic chassis, due to the differences in the transcriptional machinery in prokaryotes and eukaryotes, we sought to establish a system in *S. cerevisiae* based on the native alkane-sensing mechanism found in *Yarrowia lipolytica*.

*Y. lipolytica* is well-known for its ability to grow on hydrocarbons, such as alkanes, and the system involved in degrading alkanes has been investigated in several consecutive studies^*18-22*^. The alkane-sensing and degrading system^*18*^ consists of a cytochrome P450 enzyme, Alk1p, and at least three TFs named Yas1p, Yas2p and Yas3p (Figure 1). The *ALK1* promoter, P_*ALK1*_, has been systematically studied, resulting in the identification of an alkane-responsive region consisting of alkane responsive elements, including ARE1, which has been reported to be important for alkane-induced gene expression^*18*^. In subsequent studies^*19, 20*^, Yas1p and Yas2p were reported to bind as heterodimers to the ARE1 sequence and thereby activate transcription of *ALK1*. Furthermore, Yas3p has been reported to be involved in the transcriptional repression^*21*^ of *ALK1* when bound to Yas2p, whereas de-repression of *ALK1* is reported to occur in the presence of alkanes, through a mechanism in which Yas3p is released from Yas2p and re-localized to the endoplasmic reticulum^*21*^. However, the mechanism of how Yas3p re-localizes to the endoplasmic reticulum has not yet been completely elucidated^*22*^.

**Figure 1.**
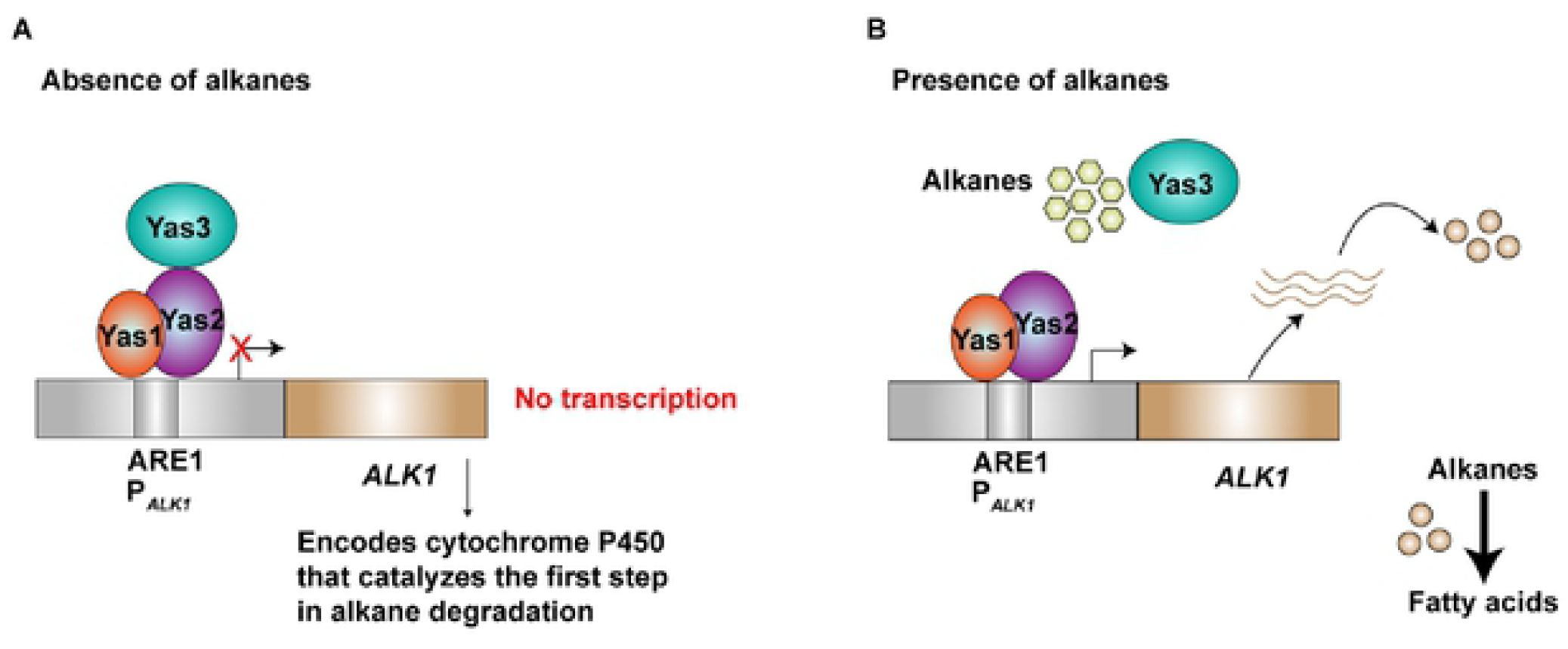
Putative alkane-sensing system in Y. lipolytica. Y. lipolytica is an alkane-assimilating yeast capable of utilizing alkanes as a carbon source. The system is, at least, based on three different transcription factors, Yas1p, Yas2p and Yas3p. A) Yas1p and Yas2p have been reported to function as an activator complex, activating expression of the ALK 1 gene encoding a cytochrome P450 enzyme that is involved in catalyzing the first step in alkane degradation whereas Yas3p acts as a repressor upon binding to Yas2p. This repression occurs in the absence of alkanes. B) On the other hand, activation occurs in the presence of alkanes as these are believed to re-localize Yas3p to the endoplasmic reticulum (not shown).

Here, we were interested in implementing the *Y. lipolytica* alkane-sensing system as a potential alkane-responsive biosensor in *S. cerevisiae*. We sought to create two alkane-responsive TF-based biosensors; one based on a synthetically modified P_*CYC1*_ promoter containing the ARE1 binding sites and one sensor system based on the native promoter P_*ALK1*_ from *Y. lipolytica*.

## RESULTS

### Establishing an alkane-responsive biosensor in *S. cerevisiae*: System based on P_*ALK1*_

When constructing TF-based biosensors in *S. cerevisiae*, the binding sites for that particular TF are normally implemented into an endogenous promoter of a desired strength^*11, 23*^. We were interested in evaluating whether it is possible to directly implement the *ALK1* promoter of *Y. lipolytica* in *S. cerevisiae*. We did this by placing P_*ALK1*_ upstream of GFP expressed from a centromeric plasmid in the background strain CEN.PK113-11C. The GFP signal, measured 6 h and 28 h after inoculation, was, however, not much higher than the signal from the same background strain carrying an empty plasmid (negative control) (Figure 2). This was not entirely unexpected as the *ALK1* promoter requires activation. Therefore, the gene encoding GFP under the control of P_*ALK1*_ was also co-expressed with the genes encoding the TFs Yas1p, Yas2p and Yas3p. As Yas1p and Yas2p together were previously reported to function as activators in *Y. lipolytica*, and co-production of Yas3p together with Yas1p and Yas2p has been reported to repress the system, we expected to achieve similar results, assuming that P_*ALK1*_ was functional in *S. cerevisiae*. However, what we observed 6 h after inoculation (glucose phase) was a clearly increased GFP signal only when all three genes encoding Yas1p, Yas2p and Yas3p were co-expressed (Figure 2A). Furthermore, from the OD measurements it could also be observed that production of Yas3p, either by itself or together with Yas1p and Yas2p, resulted in a growth defect, as seen from the OD measurements (Figure 2A). The same pattern concerning cell growth was also observed for the measurements taken 28 h after inoculation (late ethanol phase/early stationary phase), where expression of only *YAS3* or co-expression of *YAS3* together with *YAS1* and *YAS2* still resulted in a reduced OD compared to the other strains (Figure 2B). Furthermore, the GFP measurements also resulted in the same pattern as for the GFP signals observed from samples taken 6 h after inoculation (Figure 2B). However, in the measurements taken 28 h after inoculation, there was a clearer increase in the GFP signal for the strain expressing the genes encoding Yas1p and Yas2p, indicating the possibility that Yas1p and Yas2p might have some activator activity and that P_*ALK1*_ might function as promoter in *S. cerevisiae*, despite its low basal expression level (Figure 2B).

**Figure 2.**
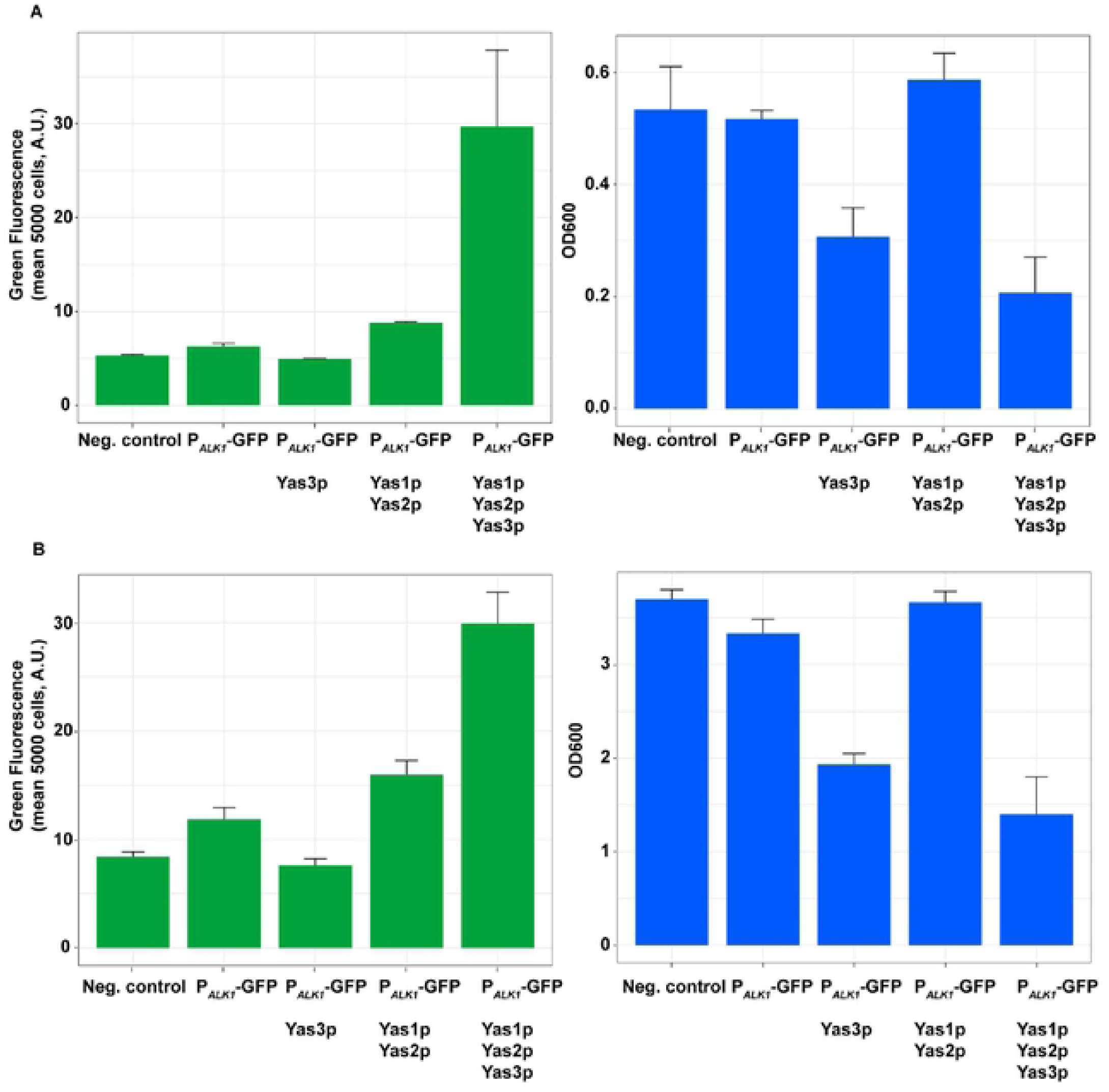
Evaluation of the alkane-sensing system in S. cerevisiae by expressing GFP from the ALK1 promoter in either the presence or the absence of the TFs Yas1p, Yas2p and Yas3p from Y. lipolytica. The promoter P_ALK1_ was placed upstream of GFP expressed from a centromeric plasmid. The genes encoding the TFs were co-expressed and placed under promoter P_PG K1_ (YAS1 ai1d YAS2) and P_TEF1_ (YAS3), respectively. Fluorescence and OD measurements were performed A) 6 h after inoculation and B) 28 h after inoculation. Strains were cultured in synthetic complete media in shake flasks. n = 3, error bar=± SD.

To further investigate GFP expression and the morphology of the cells, we evaluated the strains under the microscope 28 h after inoculation (Figure 3). As expected, the negative control did not result in any GFP expression (Figure 3A) and expression of the positive control, consisting of P_*TEF1*_ placed upstream of GFP, resulted in GFP expression (Figure 3B). However, the other strains, except for the strains containing pALK1-GFP together with the plasmid encoding Yas1p, Yas2p and Yas3p (Figure 3F), did not result in any obvious GFP signal (Figure 3C-E). Furthermore, the morphology of the strain carrying all three TFs, Yas1p, Yas2p and Yas3p, was also different from the other strains as its cells were larger and clustered together (Figure 3F). To ensure that the fluorescence signal observed in this strain was not autofluorescence, which can occur especially from dead cells, we evaluated the cell viability through staining with propidium iodide (Supporting Information, Figure S1). Cells expressing GFP did not co-stain with propidium iodide therefore indicating that the GFP signal observed from these cells was probably not due to autofluorescence. As an additional control to confirm that the observed fluorescence is not due to autofluorescence, we expressed the TFs in the absence of the GFP containing plasmid. Indeed, the observed fluorescence could only be seen for the strains expressing all three TFs together with the *ALK1* promoter coupled to GFP (Supporting Information, Figure S1). Combining the expression of the three different genes encoding the TFs, for example, co-expressing genes encoding Yas1p with Yas3p or Yas2p with Yas3p, did not result in a GFP signal, indicating again that the GFP seen is only observed when co-expressing all three TFs (Supporting Information, Figure S2). An additional observation in this experiment was that, although expressing the gene encoding Yas3p separately or together with the two other TFs resulted in a growth defect, a severe growth defect could not be observed when Yas3p was expressed with only Yas1p (Supporting Information, Figure S2).

**Figure 3.**
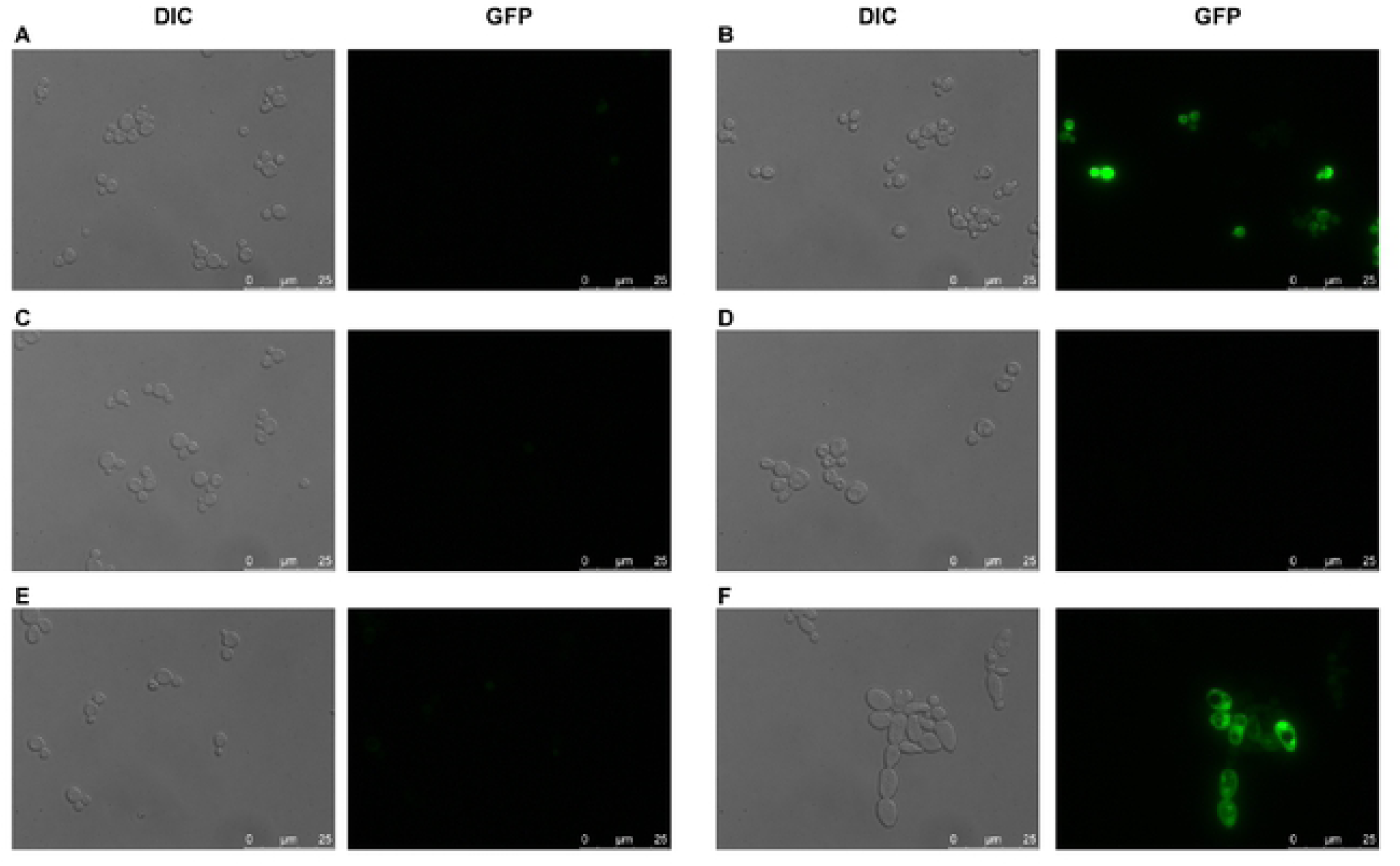
Evaluation of the strains under the microscope. **A)** Empty plasmid (negative control), B) P_TEF1_-GFP (positive control), C) P_ALK1_-GFP, D) P_ALK1_-GFP_YAS3, E) P_ALK1_-GFP_YAS1_YAS2, and F) P_ALK1_-GFP_YAS1_YAS2_YAS3. The strains were evaluated 28 h after inoculation.

### Establishing an alkane-responsive biosensor in *S. cerevisiae*: System based on P_*CYC1*_

In addition to developing the sensor system based on P_*ALK1*_, we decided to also create a system based on the endogenous promoter P_*CYC1*_. The *CYC1* promoter has been reported to be a promoter with weak/medium activity, which makes it suitable to use when developing biosensors based on activators. In fact, a study using prokaryotic activators in *S. cerevisiae* was successful in using modified versions of P_*CYC1*_ with integrated binding sites of their specific TFs^*24*^. We employed the same approach and implemented the ARE1 binding site in the *CYC1* promoter. The sequences and the positions of the binding sites can be seen in Supporting Information. The native *CYC1* promoter as well as the synthetically modified ones were placed upstream of GFP and expressed from a centromeric plasmid. The plasmids containing the different versions of P_*CYC1*_ upstream of GFP were expressed with the genes encoding the TFs Yas1p, Yas2p and Yas3p and fluorescence and OD measurements were taken 6 h and 28 h after inoculation (Figure 4). What was observed from the measurement taken 6 h after inoculation was an increased GFP signal when only expressing the activators Yas1p and Yas2p, indicating an activation. However, expressing these TFs together with Yas3p resulted in GFP signals with large variations for the triplicates, as can also be seen from the error bars (Figure 4A). The OD measurements were consistent with what has been previously observed (Figure 2), where expression of the gene encoding Yas3p resulted in reduced OD (Figure 4A). The activation pattern observed when expressing genes encoding Yas1p and Yas2p, and the growth defect observed when expressing the gene encoding Yas3p could also be observed for samples measured 28 h after inoculation (Figure 4B). However, the GFP signal for samples producing Yas3p still showed a large variation, but the overall GFP signal seemed to have increased, which gives inconclusive results regarding its repressive mechanisms or function. However, to verify that the activation observed when only expressing Yas1p and Yas2p was due to their binding to ARE1 in the *CYC1* promoter, we performed a control experiment using the unmodified P_*CYC1*_ version, to see whether a similar activation would be obtained (Supporting Information, Figure S3). Indeed, similar activation pattern could be observed when expressing the genes encoding Yas1p and Yas2p together with the plasmid carrying unmodified P_*CYC1*_-GFP.

**Figure 4.**
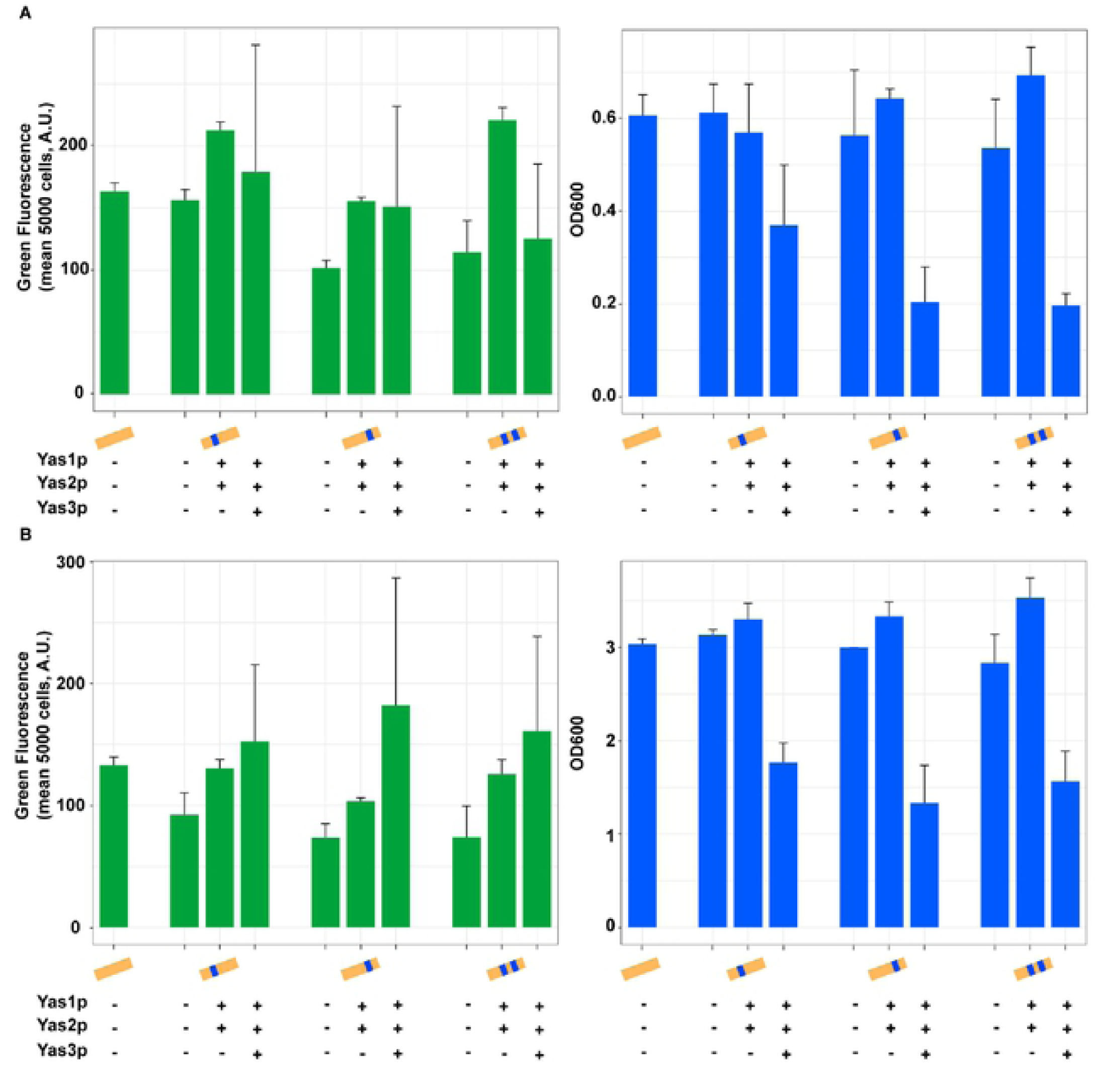
Evaluation of the alkane-sensing system in *S. cerevisiae* by expressing GFP from the *CYC1* promoter together with the genes encoding the TFs Yas1p, Yas2p and Yas3p from *Y. lipolytica*. The ARE1 binding site was integrated into the *S. cerevisiae* native yeast promoter P_*CYC1*_. The unmodified as well as the synthetically modified P_*CYC1*_ was placed upstream of *GFP* expressed from a centromeric plasmid. The gene encoding the TFs were co-expressed and placed under promoter P_*PGK1*_ (*YAS1* and *YAS2*) and P_*TEF1*_, (*YAS3*). Fluorescence and OD measurements were performed A) 6 h after inoculation and B) 28 h after inoculation. Strains were cultured in synthetic complete media in shake flasks. n = 3, error bar = ± SD.

### Evaluation of the transcription factors through fusion with GFP

To gain more insights into the production and localization patterns of Yas1p, Yas2p and Yas3p, we decided to N-terminally fuse each TF to GFP through a linker*25* consisting of the amino acids GGGS (Figure 5A). From the microscope evaluation, Yas1p seems to be localized to the nucleus as indicated by the single, concentrated, round-shaped, GFP signals (Figure 5C). On the other hand, Yas2p seems to be poorly expressed (Figure 5D) whereas Yas3p seems to be expressed, although it is difficult to identify its localization due to the scattered GFP signal across the cell with multiple foci, which could be due to targeting to unknown organelles or potentially resulting in insoluble proteins (Figure 5E). The TFs were also fused to GFP at their C-terminus to verify that any unexpected result is not due to the interaction of the GFP with the N-terminus (Supporting Information, Figure S4). Fusing GFP to the C-terminus resulted in the same GFP expression pattern as observed in Figure 5 for cells carrying Yas1p, Yas2p and Yas3p fused to GFP at their N-terminus. Furthermore, a nuclear localization signal (NLS) tag was placed at the C-terminus of the TFs Yas1p and Yas3p to verify whether the native NLS sequence was sufficient for the presumed nuclear targeting (Supporting Information, Figure S4). From the observed GFP signal, it seems that adding an additional NLS sequence did not result in a different GFP pattern, indicating that the native NLS of these TFs are sufficient (Supporting Information, Figure S4).

**Figure 5.**
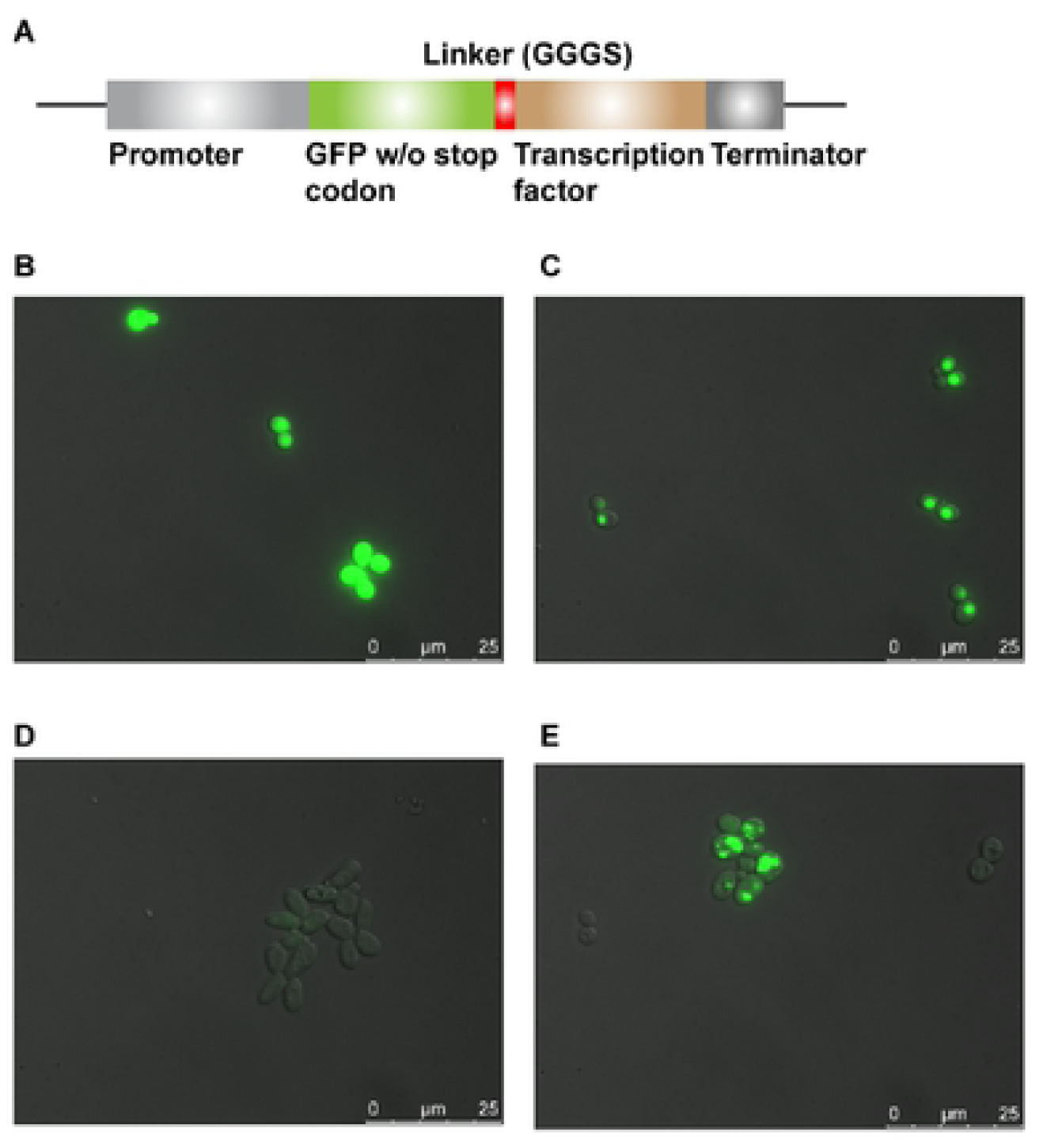
Fusion of TFs with GFP. The TFs Yas1p, Yas2p and Yas3p were N-terminally fused with GFP to evaluate their expression and localization. A) The fusion proteins were constructed through a GGGS linker. B) Positive control of GFP without fusion to any TF. C) Yas1p fused to GFP under the control of P_*PGK1*_*-*D) Yas2p fused to GFP under the control of P_*PGK1*_ E) Yas3p fused to GFP under the control of P_*TEF1*._ Samples were evaluated 6 h after inoculation.

We also evaluated these GFP-TF constructs under co-expression with other TFs (Figure 6). The fluorescence signal seen from strain carrying GFP-Yas1p together with Yas3p did not seem to disrupt Yas1p’s location to the nucleus (Figure 6A) whereas the strain carrying GFP-Yas2p together with Yas3p resulted in a brighter and clearer GFP signal (Figure 6B), which could not be observed previously (Figure 5D). Furthermore, when evaluating strains carrying Yas1p and Yas2p together with different variants of Yas3p fused with GFP, two clearly different GFP patterns were observed. The constructs containing GFP-Yas3p and Yas3p-NLS-GFP (Figure 6C and E, respectively), resulted in a clear and concentrated GFP signal located in the nucleus, whereas for Yas3p-GFP the same pattern seen previously with scattered GFP signal spread across the cell was observed (Figure 6D).

**Figure 6.**
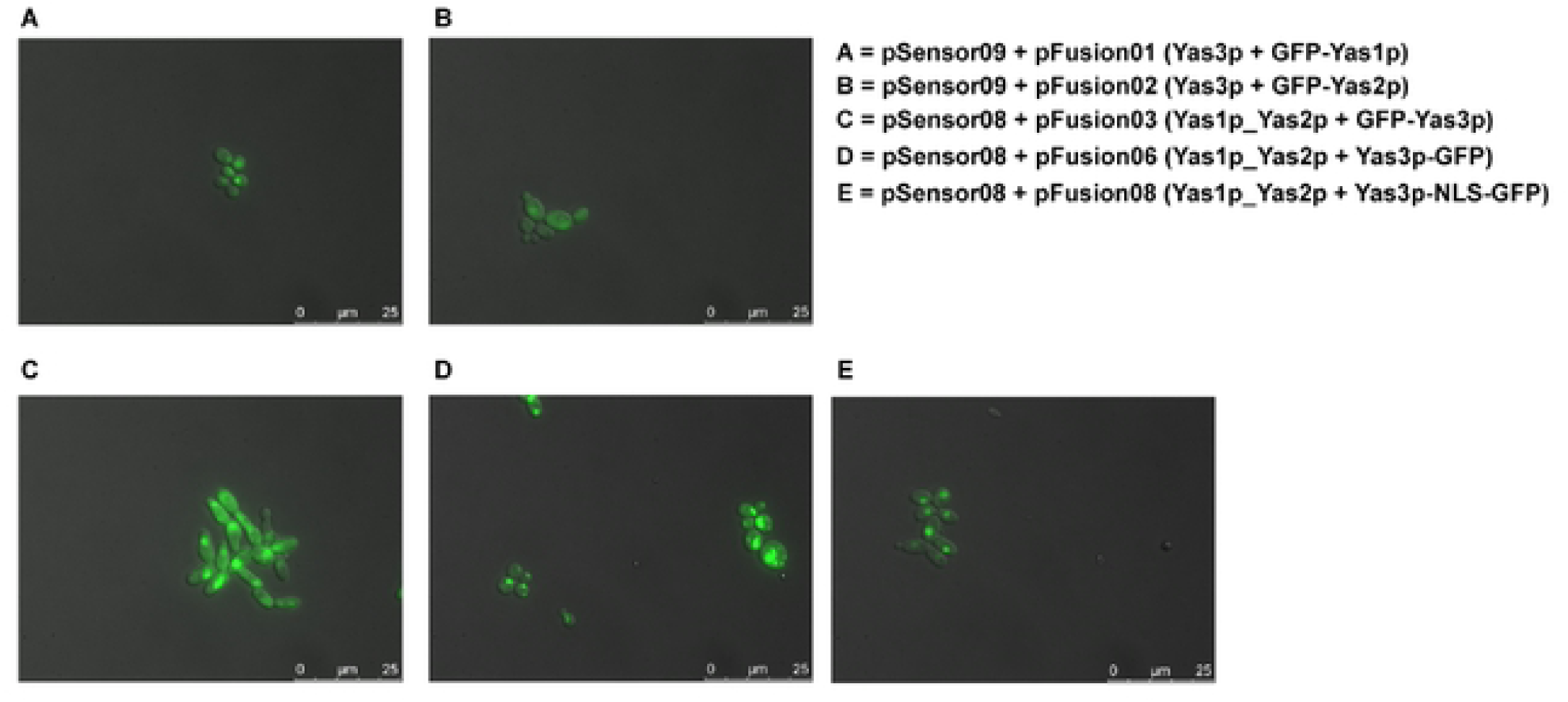
Evaluation of the GFP-tagged TFs in combination with other TFs. To better understand the interaction of the TFs with each other, co-expression of the TFs and the fusion proteins was performed. Co-expression of plasmids containing A) Yas3p with Yas1p fused to GFP, B) Yas3p with Yas2p fused, with GFP, C) Yas1p and Yas2p with GFP fused to Yas3p through its N-terminus, D) Yas1p and Yas2pwith Yas3p fused with GFP through its C-terminus, and E) Yas1p and Yas2p with Yas3p linked to an NLS tag and fused with GFP through its C-terminus. Samples were evaluated 6 h after inoculation.

## DISCUSSION

The use of metabolite biosensors, such as an alkane-responsive biosensor, as high-throughput screening tools can speed up the search for high-performing cell factories. Here, we investigate whether an alkane-responsive biosensor based on the three TFs Yas1p, Yas2p and Yas3p from the native alkane-sensing system in *Y. lipolytica* can be transferred to *S. cerevisiae*. Two putative alkane-responsive, TF-based, biosensors were developed and evaluated; one based on the native *S. cerevisiae* promoter P_*CYC1*_ and one based on the native *Y. lipolytica* promoter P_*ALK1*_.

Although it is rare to use heterologous promoters from a distantly related species in a given host, P_*TEF1*_ from *Ashbya gossypii* has successfully been used in *S. cerevisiae*^*26*^. The alkane-sensing system that naturally exists in *Y. lipolytica* is based on the P_*ALK1*_ promoter and the TFs Yas1p, Yas2p and Yas3p. Although *Y. lipolytica* does not belong to the Saccharomycetaceae family like *S. cerevisiae* and *A. gossypii, S. cerevisiae* and *Y. lipolytica* are both Ascomycota of the order Saccharomycetales. Therefore, we decided to directly implement the P_*ALK1*_ promoter in *S. cerevisiae* and evaluate whether this implementation would be feasible for developing an alkane-responsive sensor system based on the three TFs.

From our studies, it is not clear whether P_*ALK1*_ activates gene expression in *S. cerevisiae*. In a previous study*27* it has been shown that the expression level of *ALK1* gene in nitrogen limiting conditions is comparable to the average expression of all other genes, which might indicate that there is some basal activity of the *ALK1* promoter in its native host. We did, however, not observe any basal activity above the background level. Furthermore, the activating function of Yas1p and Yas2p was inconclusive, since they seemed to only mildly activate expression as seen from the increased fluorescence signal observed when employing flow cytometry, whereas no signal could be detected when evaluating the strains using microscopy.

According to the alkane-sensing and degrading system reported in *Y. lipolytica*, expression of all three genes encoding the TFs Yas1p, Yas2p and Yas3p was expected to result in repression as Yas3p has been shown to bind to Yas2p and repress transcription. In contrast to what has been reported, expressing all three genes encoding the TFs resulted in increased GFP production under the control of P_*ALK1*_, which we also observed through microscopy. Despite testing various combinations of the TFs, the fluorescence signal was only observed when expressing all three genes encoding Yas1p, Yas2p and Yas3p together. Although Yas3p has been reported to function mainly as a repressor, its mechanism is not yet fully understood as it has also been reported to have a positive regulatory function^*21, 22*^.

We also observed growth defect when expressing all genes encoding the TFs together or when expressing only the gene encoding Yas3p. The observed growth defect is probably due to the relatively high expression levels in this study, using P_*PGK1*_ for expressing Yas1p and Yas2p and using P_*TEF1*_ for expressing Yas3p. To find out whether the observed growth defect is due to the high expression levels, we performed an additional experiment (data not shown) where the gene encoding Yas3p was placed under the control of P_*OPI1*_ and P_*REV1*_, which were chosen since Yas3p might play a similar role as Opi1p in *S. cerevisiae* and since P_*REV1*_ is reported to be a promoter with very weak strength. Although the growth defect was less severe when using P_*OPI1*_ and P_*REV1*_ than in the experiment of P_*TEF1*_-based expression, the putative alkane-sensing system expressed with GFP under the control of P_*CYC1*_ containing ARE1 binding sites was still not showing any signs of repression as measured by the GFP signal (data not shown). This observation indicates that the observed phenotype is not only due to the overexpression of Yas3p, but that there are other factors that prevent the system from functioning.

As it is not clear whether P_*ALK1*_ functions properly in *S. cerevisiae*, we decided to use a native *S. cerevisiae* promoter, the *CYC1* promoter, to evaluate the sensor system. Using P_*CYC1*_, activation could be observed when co-expressing the genes encoding Yas1p and Yas2p. However, this activation was also observed when expressing the activator complex with the unmodified P_*CYC1*_ construct, which did not contain any ARE1 binding sites. After a closer examination, a part of the motif (CTTGTGN_X_**CATGTG**), to which Yas1p and Yas2p are reported to bind^*20*^, was also found to be present in the P_*CYC1*_ sequence (CATGTG). This motif has also been recognized by the native Ino2p-Ino4p TF complex. A question that remains to be answered is whether the activation observed in these strains is due to Yas1p and Yas2p binding to the ARE1 element and thereby activating transcription or whether these two TFs activate the promoter by binding to a different site in its native sequence.

To obtain a better understanding of how the TFs are expressed in *S. cerevisiae*, we tagged each TF to GFP. Despite both Yas1p and Yas2p being produced from the promoter P_*PGK1*_, production of Yas1p and its localization to the nucleus was clear whereas Yas2p seemed to be weakly expressed, making it difficult to evaluate its production and localization. The challenge of evaluating the production of Yas2p through fusion with GFP was also encountered in a previous study^*20*^ in *Y. lipolytica*; this behavior can potentially be due to Yas2p being an unstable protein in the absence of Yas1p and/or Yas3p. In a subsequent study^*21*^ the authors were, however, successful in detecting Yas2p fused with GFP in the nucleus. We did not evaluate the expression of the gene encoding Yas2p together with the gene encoding Yas1p, but from the co-expression of the genes encoding Yas2p and Yas3p, an observation of a clearer GFP signal indicates that the stability of Yas2p might be enhanced by Yas3p. Furthermore, in our study, Yas3p seemed to be produced although its localization could not be clearly elucidated as the GFP was observed to be spread throughout the whole cell as foci of different sizes. It seems that Yas3p has some interdependence with the TFs Yas1p and Yas2p as it was observed to have less dispersed GFP signal in some of the cases when these were co-expressed together. Binding to Yas1p and Yas2p may have changed the pattern of Yas3p by channeling it to a different localization. If the foci indicate insoluble protein, a co-expression could have influenced the folding process.

It has been suggested that the alkane-sensing system based on Yas1p, Yas2p and Yas3p shares similarities with the system regulating lipid biosynthesis in *S. cerevisiae*, which is based on the heterodimeric Ino2p-Ino4p activator complex and the transcription factor Opi1p. The activators Ino2p-Ino4p are global regulators reported to regulate the expression of genes involved in lipid metabolism as well as genes unrelated to phospholipid metabolism^*28-30*^. The effector molecules of this system are commonly inositol and choline, which regulate genes encoding phospholipid-synthesizing enzymes. When inositol and choline are scarce, Ino2p-Ino4p activates the expression of genes involved in phospholipid synthesis whereas when inositol and choline are available, Opi1p represses this system. Similar to the mechanism for Yas3p, which is reported to change its location to the endoplasmic reticulum in the presence of alkanes, Opi1p sequesters to the endoplasmic reticulum membrane in the absence of inositol. Furthermore, when comparing the amino acid sequences of Yas3p and Opi1p, some similarities between these two exists, including the presence of a leucine zipper domain, similarity in an uncharacterized domain and similarities in the activator interaction domain^*21*^. Despite the similarities shared between these systems, the system in *Y. lipolytica* has been reported not to be sensitive to inositol. The roles of Yas1p-Yas2p and Yas3p may, however, be as complex as that of the Ino2-Ino4p and Opi1 system, particularly as there is still much to be understood regarding the mechanisms of Opi1p and Yas3p.

In summary, our findings suggest that before an alkane-responsive biosensor can be developed based on the alkane-sensing system in *Y. lipolytica*, it is necessary to gain a deeper understanding of the TFs and their mechanisms. Normally, TF-based biosensors are constructed using a well-characterized, single TF, whereas a biosensor based on three TFs adds a greater complexity to the system and makes fine-tuning of it a challenging task. When expressing the genes encoding the TFs Yas1p, Yas2p and Yas3p, we encountered problems in achieving high expression levels, growth defects and problems in obtaining clear data on the TFs cellular localization. We do not exclude the fact that such a sensor can be constructed based on this alkane-sensing system. We believe, however, that such a system is not mature to be translated into a biosensor in *S. cerevisiae* until a better understanding of these TFs has been achieved and potential additional components involved in the mechanisms have been identified. Our study demonstrates an example of the challenges that occurs when developing a metabolite biosensor, and the need to possess a well-characterized regulatory system before transforming such a system to an orthologous biosensor that can be applied in different hosts.

## MATERIAL AND METHODS

### Chemicals and Reagents

All primers were synthesized at Eurofins. Restriction enzymes, DNA gel extraction- and plasmid purification kits were purchased from Thermo Fischer Scientific. The Gibson Assembly® Master mix, purchased from New England Biolabs, was used for plasmid construction. Phusion High-Fidelity DNA Polymerase (ThermoFisher Scientific) was routinely used for PCR amplification. All reagents used for media preparation were purchased from Merck Millipore unless otherwise noted. Propidium iodide (PI) was purchased from Invitrogen.

### Strains

*S. cerevisiae* strain CEN.PK113-11C (*MAT*a*SUC2 MAL2-8*^*c*^ *his3*Δ*1 ura3-52*)^*31*^ was used as the background strain for all experiments. For standard cloning procedures competent *Escherichia coli* cells, DH5α, were routinely used. Genomic DNA of CEN.PK113-11C was used as a template, unless otherwise noted, when amplifying endogenous promoters or terminators. Genomic DNA of *Y. lipolytica*, strain FKP391^*32*^, was used for amplifying promoter P_*ALK1*_.

### Media

Complete medium, YPD, containing 10 g/L yeast extract, 20 g/L casein peptone and 20 g/L glucose, was used when preparing yeast competent cells. For selection of yeast transformants carrying *URA3-* and *HIS3*-based plasmids, synthetic complete media plates without uracil and histidine (SC-URA-HIS) containing 6.7 g/L yeast nitrogen base (YNB) without amino acids (Formedium), 0.77 g/L complete supplement mixture without uracil and histidine (CSM-URA-HIS, Formedium), 20 g/L agar and 20 g/L glucose were used.

For fluorescence analysis, yeast strains were cultured in synthetic complete media containing 6.7 g/L yeast nitrogen base without amino acids, 0.77 g/L CSM-HIS-URA, and 20 g/L glucose. For culturing *E. coli* cells, Luria-Bertani (LB) broth supplemented with 100 mg/L ampicillin was used.

### Plasmid- and strain construction

*S. cerevisiae* strain CEN.PK113-11C was transformed with the following plasmids using the lithium acetate method^*33*^. All plasmids were verified through restriction digestion analysis and sequencing at Eurofins.

Plasmid p413TEF1-GFP was constructed by amplifying the backbone plasmid p413TEF1 using primer pair pYDA01/02 and primer pair pYDA03/04 to amplify the GFP gene. Similarly, plasmid p413CYC1 was constructed by using primer pair pYDA01/05 to amplify backbone plasmid p413TEF1 and primer pair pYDA06/07 to amplify P_*CYC1*_ from genomic DNA CEN.PK113-11C. Plasmid p413CYC1-GFP was constructed by using p413TEF1-GFP as a backbone plasmid and primer pair pYDA05/08 to linearize it and assemble it with P_*CYC1*_, which was amplified using primer pair pYDA06/09. The same, linearized backbone plasmid p413TEF1-GFP, was assembled with P_*ALK1*_, which was amplified from *Y. lipolytica* strain FKP391 using primer pair pYDA10/11, resulting in plasmid pALK1-GFP.

Plasmids pSensor01-03 were constructed using primer pair pYDA12/09 to amplify part of P_*CYC1*_ with the ARE1 BS included (purchased as gBlock fragments from Integrated DNA Technologies) and primer pair pYDA13/08 to amplify the backbone p413CYC1-GFP such that the P_*CYC1*_ part included in the gBlock fragment was excluded from the backbone. To construct the plasmids, pSensor04-08, pSensor01-03, p413CYC1 and pALK1-GFP were digested with PstI and assembled with cassette P_*PGK1*_-*YAS1*-T_*ADH1*_ (Genscript_001) amplified using primers pYDA14/15, cassette *YAS2*-T_*PDC6*_ (Genscript_002) amplified using primer pair pYDA16/17 and P_*PGK1*_ overlapping with *YAS2*-T_*PDC6*_ using primer pair pYDA18/19. pSensor09 was constructed by amplifying the backbone plasmid p416TEF1 using primer pair pYDA05/20 and primer pair pYDA21/22 to amplify P_*TEF1*_-*YAS3*-T_*CYC1*_ (Genscript_003).

pFusion01 was constructed by amplifying p413TEF1 using primer pair pYDA05/20, P_*PGK1*_ from genomic DNA using primer pair pYDA23/24, GFP (without stop codon, with linker GGGS^*25*^) using primer pair pYDA25/26, *YAS1* and T_*ADH1*_ separately from Genscript 001 using primer pair pYDA27/28 and pYDA29/30. pFusion02 was constructed similarly with minor changes; GFP was amplified using primer pair pYDA25/31 and to amplify *YAS2* primer pair pYDA32/33 were used. pFusion04-05 were constructed similarly with minor changes in primer pairs. To construct pFusion04, pYDA39/40 amplifying *YAS1* (without stop codon, with linker GGGS^*25*^) and primer pair pYDA41/42 amplifying *GFP* were used. For constructing pFusion05, pYDA43/44 were used to amplify *YAS2* (without stop codon, with linker GGGS^*25*^) and pYDA45/46 to amplify *GFP*, and T_*PDC6*_ using primer pair pYDA53/33. pFusion07 is similar to pFusion04 with the difference that pFusion07 has an NLS tag (PKKKRKV) included in the C-terminus of *YAS1*. pFusion07 was consequently constructed similarly to pFusion04 using primer pair pYDA39/50 to amplify *YAS1* and primer pair pYDA51/42 to amplify GFP. pFusion03 was constructed using primer pair pYDA05/34 to amplify backbone plasmid pSensor09. Primer pair pYDA35/36 was used to amplify *GFP* (without stop codon, with linker GGGS) and primer pair pYDA37/38 to amplify P_*TEF1*_. Similarly, pFusion06 was constructed using pSensor09 as a backbone plasmid, which was amplified using primer pair pYDA01/47 and subsequently assembled with *GFP*, which was amplified using primer pair pYDA48/49. pFusion08, similar to pFusion06, with the difference that pFusion08 contains an NLS tag (PKKKRKV) in the C-terminus of *YAS3*, was constructed similarly to pFusion06 with the difference in primer pair amplifying *GFP*. Here, primer pair pYDA48/52 was used to amplify *GFP* and assemble it with the linearized background plasmid pSensor09, which was amplified using pYDA01/47.

pSensor10 was constructed by digesting plasmid pALK1-GFP with PstI and assemble it with P_*PGK1*_-*YAS1*-T_*ADH1*_, which was amplified from fragment Genscript 001 using primer pair pYDA55/56. Similarly, pSensor11 was constructed by amplifying fragment *YAS2*-T_*PDC6*_ using primer pair pYDA54/17 and P_*PGK1*_ using primer pair pYDA18/19, which were assembled with the linearized pALK1-GFP plasmid.

The amplified backbone plasmids pSensor09 using primer pair pYDA05/34 was assembled with P_*OPI1*_, amplified using primer pair pYDA57/58, which resulted in plasmid pSensor12. Similarly, plasmid pSensor13 was constructed by assembling linearized pSensor09 with P_*REV1*_, which was amplified using primer pair pYDA59/60.

**Table 1.**
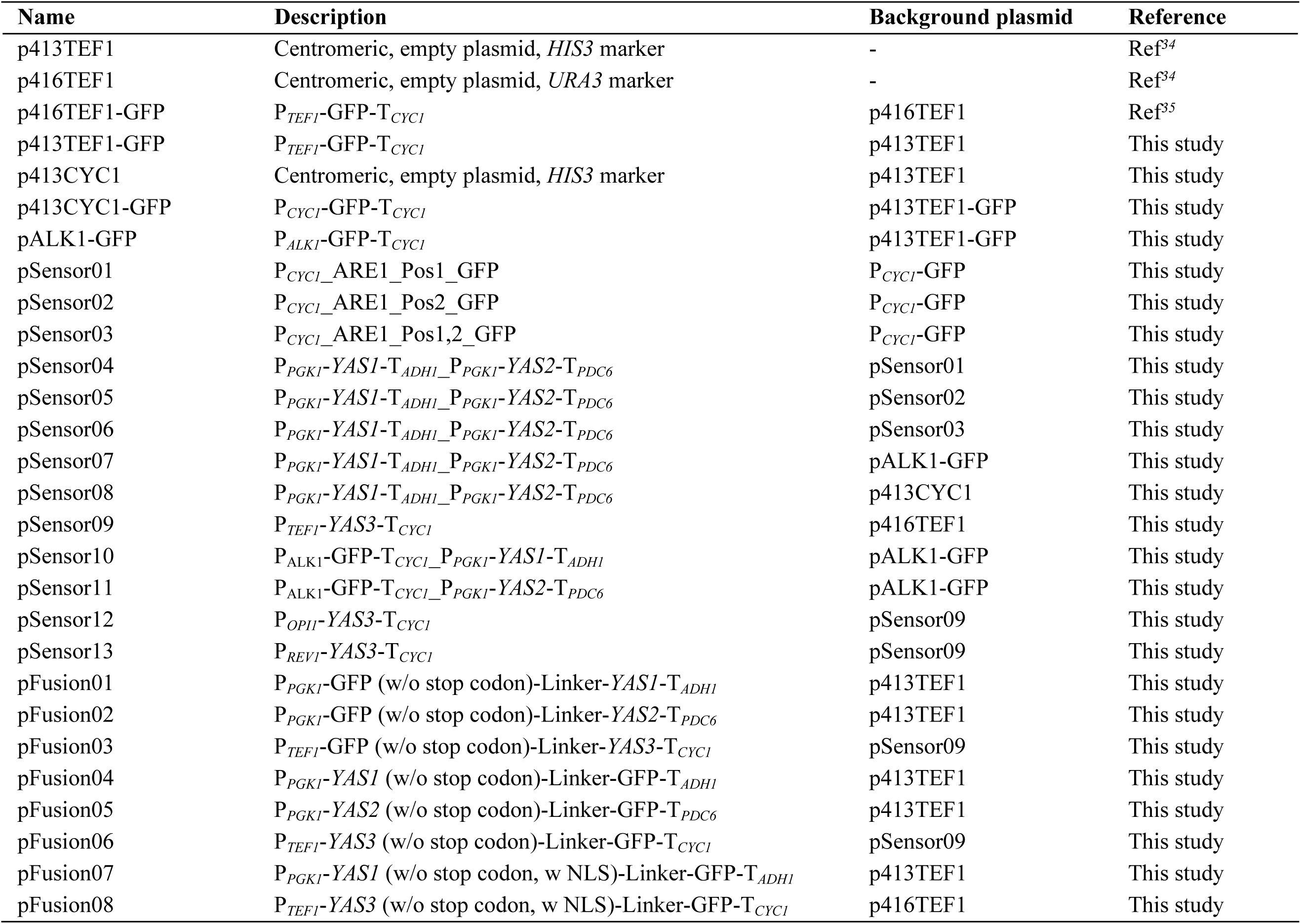
Plasmids used in this study

### Flow cytometry measurements

For flow cytometry analysis, Guava easyCyte 8HT system (Merck Millipore, Billerica, MA, USA) with a blue laser (405 nm) and green fluorescent filter (520/30 nm) was used. Cultures were inoculated at an OD_600_ of 0.1 in 10 mL media in 100 mL shake flasks. Samples were analyzed 6-8 h and 24-26 h after inoculation of an OD_600_ 0.1. All samples were, prior to analysis, diluted in water to an OD_600_ of 0.02 in a final volume of 200 μl, and 5000 cells were measured.

### Propidium iodide (PI) staining

To evaluate the viability of some of the strains, cells were resuspended with 1 mL phosphate-buffered saline (PBS) with 0.5 μl of 1 mg/mL PI. Samples were incubated in dark for 20 min, followed by centrifugation at 1000 g for 4 min. The supernatant was discarded, and the cells were resuspended in 10 μl PBS and evaluated under microscope. The positive control consisting of dead cells were obtained by boiling the WT cells in 100°C for 20 min and following the same protocol as described above.

## ASSOCIATED CONTENT

## Supporting Information

Cell viability evaluation through propidium iodide (PI) staining (Figure S1); Evaluation of Yas1p, Yas2p and Yas3p with the *Yarrowia lipolytica* promoter *ALK1* (Figure S2); Evaluation of Yas1p and Yas2p together with the endogenous promoter *CYC1* as well as a synthetically modified version with ARE1 BSs (Figure S3); Fusion of TFs Yas1p, Yas2p and Yas3p with GFP (Figure S4); primers (Table S1) and sequences.

## ACKNOWLEDGMENTS

This work was funded by the Novo Nordisk Foundation (grant no. NNF10CC1016517) and ÅForsk, which is gratefully acknowledged. The authors would also like to acknowledge Xin Chen for helpful discussions, John Hellgren for providing genomic DNA of *Y. lipolytica* FKP391 and Moltas Olofsgård, Marcus Baaz, Martin Tabell, Alex Hedin, Elin Johansson and Malin Stenbom for great assistance with some of the experiments.

## AUTHOR CONTRIBUTIONS

Y.D., V.S., and F.D., designed the research; Y.D., performed all the experiments; C.S., assisted in an experiment, Y.D., analyzed the results. All authors discussed the results; Y.D., prepared the original draft and F.D., and V.S., helped reviewing and editing the draft.

## AUTHOR INFORMATION

## Notes

The authors declare no competing financial interest.

